# Reliability of self-reported catch and effort data via smartphone applications in a multi-species recreational fishery

**DOI:** 10.1101/2025.01.31.635761

**Authors:** Bernat Morro, Inmaculada Riera, Antoni Mira, Clara Mecinas, Antoni Mª Grau, Josep Alós

**Affiliations:** Mediterranean Institute for Advanced Studies, IMEDEA (CSIC–UIB), Esporles, Mallorca (Spain); Direcció General de Pesca - Government of the Balearic Islands, Palma, Mallorca (Spain); Tecnologías y Servicios Agrarios, S.A., S.M.E., M.P - Tragsatec, Palma, Mallorca (Spain)

**Keywords:** Captures per Unit of Effort, CPUE, creel survey, fishing modalities, MPA, temporal variability

## Abstract

The spatial and temporal heterogeneity of recreational anglers and the challenges associated with monitoring their activities often complicate their effective management. Angler smartphone applications (apps) offer a promising digital tool for self-reporting fishing effort (E) and captures per unit of effort (CPUE). However, despite their growing use for data collection in recreational fisheries, the existing literature on their performance remains limited, raising concerns about potential biases in the collected data. Since 2019, self-reporting of E and CPUE has been mandatory for recreational fishing trips within the network of partially-protected Marine Protected Areas (MPAs) in the Balearic Islands (Spain). A regional-scale App (“Diari de Pesca Recreativa”) was developed to streamline data collection. This study aimed to evaluate the App’s performance in reporting recreational fisheries data over a six-year period. Data obtained via the App (3,672 trip self-reports) were compared to data collected through a traditional method (360 on-site creel surveys). Estimates of E and CPUE were compared across datasets overall, as well as by month, fishing modality, MPA, and for key target species. Significant differences were observed for E (hours · angler · trip, p < 0.001) and CPUE (captures· E^-1^, p = 0.001) across datasets. However, when stratified by species, month, fishing modality, and MPA, most groups showed no statistically significant differences in E and CPUE estimates. Data from the App tended to overestimate E compared to creel surveys, likely due to inaccuracies in self-reported trip durations. Conversely, creel surveys appeared to underestimate E, reflecting intrinsic biases in trip duration collection. Assuming that the true values of E and CPUE lie between the estimates derived from the App and creel surveys, integrating both methods is recommended to improve the accuracy and reliability of data. The App not only generates a higher volume of trip data but also provides a user-friendly platform for self-reporting. Moreover, it digitizes data collection, enabling automation and advanced analytics for fisheries monitoring and management of recreational fisheries.

## 1. Introduction

Recreational fishing has direct, significant ecological and socio-economic implications in the dynamics of ecosystems (Arlinghaus et al., 2019). With millions of anglers participating worldwide, recreational fishing transformed into a multifaceted socio-economic and ecological phenomenon (Hyder et al., 2018). While recreational fisheries generate substantial benefits for people (McManus et al., 2011), when not properly managed can induce significant ecological effects, including significant modification of fish abundance (Alós et al., 2012; Lewin et al., 2019; Schroeder & Love, 2002), changes in fish behaviour (Alós et al., 2016; Monk et al., 2021; Sutter et al., 2012), decreasing average body size (Alós & Arlinghaus, 2013; Lewin et al., 2019; Monk et al., 2021), or modifying the shape of fish (Alós et al., 2014). These effects arise from both the direct harvesting of biomass and the selective nature of recreational fishing, which disproportionately targets phenotypes that are more vulnerable (Lennox et al., 2017). To mitigate these potential negative consequences, science-informed fisheries management strategies are essential (Post et al., 2002). The adoption of novel technologies is particularly crucial in achieving this goal.

Fisheries science and management have traditionally relied on cost- and time-intensive methods to collect effort (E) and captures per unit of effort (CPUE) data from recreational fisheries, primarily due to the high participation rates and the heterogeneous spatial and temporal distribution of fishing E. The most common methods for data collection include diary log-books (Jansen et al., 2013; Stephens & MacCall, 2004), creel survey (Nieman et al., 2021; Tucker et al., 2024), telephone call surveys (Arlinghaus & Mehner, 2003; Teixeira et al., 2016), and more recently cameras (Hartill et al., 2016; Signaroli et al., 2024), among others. In today’s highly interconnected digital era, digital tools hold significant potential to transform data collection in recreational fisheries (Brownscombe et al., 2019; Lennox et al., 2022). Among these tools are recreational fishing smartphone applications (apps), which attract users by providing functionalities such as logging catches and locations, and sharing fishing experiences on social platforms (Skov et al., 2021). These apps also facilitate collaboration by enabling recreational anglers to actively participate as citizen scientists, offering an accessible avenue for contributing to scientific research and fisheries management.

The number of studies comparing the performance of conventional methods and apps to assess their reliability and effectiveness remains limited. However, when compared to traditional recreational fishing survey methods, app-based data have been shown to accurately reflect regional fishing patterns and catch rates (Johnston et al., 2022) or provide comparable catch rates despite significant spatial variability in the fishing effort (Jiorle et al., 2016). However, they are not inherently suitable for estimating fish abundance or community composition as important differences from apps to conventional methods were highlighted in species-specific catch (Johnston et al., 2022; Papenfuss et al., 2015). Surveys show a positive interest in app data among researchers and managers but advise caution due to the intrinsic bias of the data (Skov et al., 2021; Venturelli et al., 2017). App users are typically younger, more specialized, and more avid anglers, often exhibiting higher catch rates compared to non-users (Griffiths et al., 2013; Gundelund et al., 2020; Trinnie & Ryan, 2024). This skew toward dedicated anglers may inflate certain metrics, such as CPUE, necessitating bias corrections to ensure the data’s reliability for management purposes (Gundelund et al., 2020; Jiorle et al., 2016). Despite inherent challenges, fishing apps present valuable opportunities for continuous monitoring, significantly enhancing both the spatial and temporal scope of data collection. These apps allow for the tracking of angler behaviour and provide a platform for innovative research applications (Papenfuss et al., 2015). By expanding the geographic and temporal coverage of data, angler apps enable the visualization of broad regional fishing patterns and improve data resolution. However, the adoption of these digital tools also introduces specific biases and challenges that must be carefully considered to optimize their effectiveness in recreational fisheries monitoring.

The objective of this study is to evaluate the performance of an angler app in contributing to the collection of fishing E and CPUE data over a six-year period, within a multi-species recreational fishery. This study is conducted within a framework that includes infrastructure to manage data collection and mitigate biases. We compared app-collected data (self-reported by anglers) to data from coastguard inspections of recreational fishing vessels (on-site creel surveys), aiming to identify potential sources of bias in each data source and to compare their estimates of E and CPUE. The analysis was applied to a controlled recreational fishery within the 12 marine protected areas (MPAs) of the Balearic Islands, spanning all 12 months and considering 8 different fishing modalities. This study represents one of the first efforts to provide evidence that smartphone apps can support data collection in spatially structured, multi-species recreational fisheries.

## 2. Methods

### 2.1. Case study: The multi-species recreational fishery in Balearic MPAs

Recreational fishing is very popular in the Balearic Islands (NW Mediterranean). In 2005, there was an estimated 5%–10% of the active population taking part (70,000–100,000 fishers, Morales-Nin et al., 2005). Most of them (63%) used recreational boats, representing around 20,000 vessels (Morales-Nin et al., 2015). Recreational fishing harvest estimates accounts for up to 25% of the total official landings in the Balearic archipelago (Morales-Nin et al., 2005). In 2024, approximately 12,000 individual and 30,000 boat-based recreational fishing licenses are active in the Balearic waters. Due to the high participation rates and the harvest-oriented nature of this fishery, local managers have implemented several management measures to ensure its ecological and socio-economic sustainability. These measures include harvest regulations such as seasonal closures, minimum size limits, gear restrictions, and the establishment of a network of partially protected MPAs (Grau, 2008).

Within this network of MPAs, certain types of recreational fishing are permitted under specific regulations designed to balance ecological conservation with recreational use(Alós & Arlinghaus, 2013). Since 2019, new legislation has been implemented in the Balearic Islands (Spain), specifically affecting recreational fishing within its 12 MPAs (BOE Decree 67/2019, of March 19, 2019, available at https://www.boe.es/boe/dias/2019/03/19/pdfs/BOE-A-2019-3911.pdf). Under these regulations, boat anglers are required to report detailed information for each fishing trip, including the fishing modality used, the number of captures from a list of 80 species (7 of which currently lack data), covering 20 families (77 fish species and 3 cephalopods), as well as additional data necessary to calculate fishing E) and CPUE of the trip. Catch self-report is focused on the harvested fraction of the catch. To facilitate these self-reports of the fishing trip, an angler smartphone application was developed and managed by the local governing and administrative authorities (see description of the App below).

Since 2022 fishing within all Balearic MPAs requires of a boat-issued fishing authorization for all boats, issued by the same local governing and administrative body, which is free and lasts for 3 years. Recreational fishing boats wishing to fish within an MPA must first obtain specific authorization for the specific MPA, comply with the associated regulations, and submit a self-report detailing their fishing trip via the App every time they fish in the MPA. MPAs are actively enforced by the governing body’s coastguards, and non-compliance with regulations results in sanctions. The mandatory nature of reporting, coupled with the extensive control over data collection, presents a unique opportunity for recreational fisheries science to evaluate the potential of using angler apps for systematic data collection. The different MPAs were incorporated into the App sequentially, presented here with the year of incorporation in parenthesis: “Llevant de Mallorca” (2018), “Sa Dragonera” (2019), “Illes del Toro i Malgrats” (2019), “Costa Nord-Est d’Eivissa-Tagomago” (2019), “Punta de sa Creu” (2019), “Illa de l’Aire” (2019), “Badia de Palma” (2020), “Migjorn de Mallorca” (2020), “Freus d’Eivissa i Formentera” (2021), “Nord de Menorca” (2022), “Ses Bledes” (2023), and “Vedrà-Vedranell” (2023). The locations of these MPAs are shown in Figure 1. Each MPA has specific characteristics and fishing rules that can be found here: www.caib.es/sites/reservesmarines.

**Figure 1.**
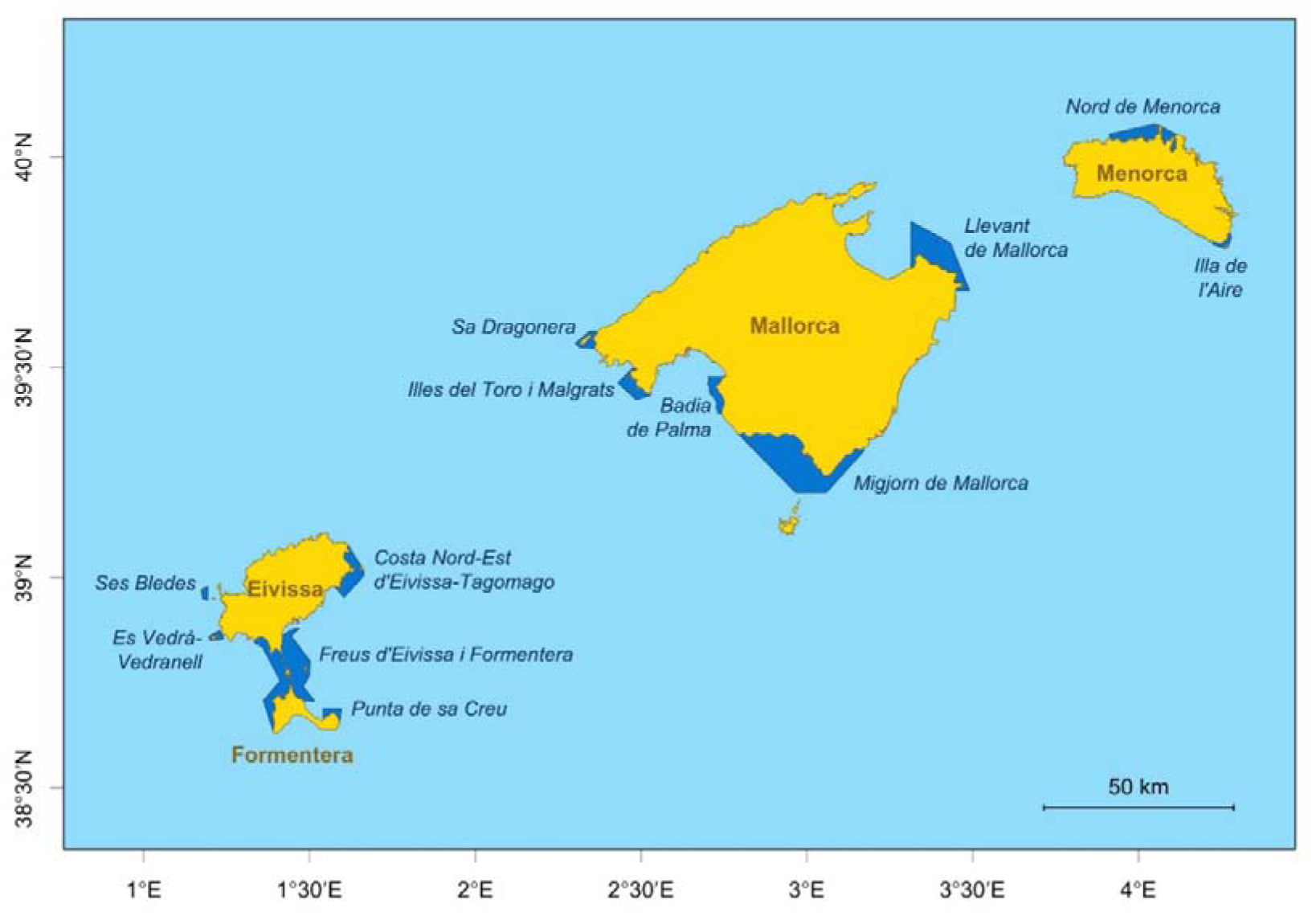
Map showing the location and delimitation of the Marine Protected Areas of the Balearic Islands, where self-reporting of effort and catch per trip is mandatory for recreational fishing.

The boat-based recreational fishery in the Balearic Islands is characterized by its multimodal nature, targeting a diverse array of species (Morales-Nin et al., 2005). This study identifies and represents eight distinct fishing modalities conducted from boats, each of which is described in the context of fishing practices in the Balearic Islands. These modalities include:

i. Trolling fishing: While the boat is cruising at slow velocities, artificial or natural lures iare dragged on the surface or upper meters of the water column, aiming for *Seriola dumerilii*, *Coryphaena hippurus*, *Sphyraena viridensis* or *Euthynnus alletteratus,* among others pelagic species.
ii. Bottom trolling fishing: While the boat is moving, a live bait (usually a squid or bait fish) is dragged close to the bottom at slow velocities, aiming for large-bodied individuals of *Seriola dumerilii*, *Dentex dentex*, or *Epinephelus marginatus*, among others. This modality is not allowed in MPAs “Punta de sa Creu” and “Illa de l’Aire”.
iii. Spinning: Done statically, a weighed jig is lowered to a depth and pulled to the surface while imitating the movement of a live fish. It aims for fish species from both Trolling fishing and Bottom trolling fishing. This modality is not allowed in MPAs “Punta de sa Creu”, “Illa de l’Aire”, “Llevant de Mallorca”, and “Costa Nord-Est d’Eivissa-Tagomago”.
iv. Squid fishing: Fishing for *Loligo vulgaris* can be done statically or moving, using specialized lures called squid jigs or egi jig. See a detailed description of the fishery in Cabanellas-Reboredo et al. (2017).
v. Razorfish fishing: Fishing for *Xyrichtys novacula* is done statically over sand patches using small hooks and normally small live bait like shrimp or worms. Other non-target species are often caught, like *Trachinus draco, Bothus podas or Balistes carolinensis.* See a detailed description of the fishery in Alós et al. (2016).
vi. Shallow fishing: A popular bottom fishing method, done statically near the shore over *Posidonia oceanica* and it mostly catches *Serranus scriba*, *Diplodus annularis* and *Coris julis*. See a detailed description of the fishery in Alós et al. (2009)
vii. Deep fishing: Another bottom fishing method, done statically and deeper than Shallow fishing that mainly aims for *Serranus cabrilla* but catches a wide variety of other species, like *Diplodus vulgaris*. Adult *Pagrus pagrus*and *Scorpaena scrofa* are the most coveted prize fish in this modality. Spearfishing: Freediving with a speargun, with a wide variety of target species depending on the spearfisher’s interests and ability. This modality is only allowed in two MPAs: “Badia de Palma” and “Migjorn de Mallorca“. See a detailed description of the fishery in Coll et al. (2004).

### 2.2. App description and App data

The App, called “Diari de Pesca Recreativa” (Angler’s Diary in Catalan), was launched in September 2018, initially with 156 users. It is a modular Java application deployed on a JBoss Enterprise Application Platform 7 server, using JDK 11 and integrated with centralized authentication via KeyCloak. The system consists of two main modules: (1) a Backoffice developed with Vue-js and JAXRS Rest services, accessing data through JPA (Hibernate) connected to a PostgreSQL database; and (2) a Frontoffice, a Vue-js Progressive Web Application (PWA) publicly accessible via the internet, which utilizes a secure Rest API (JAX-RS) specifically designed for data query and update operations. Both the Backoffice and Frontoffice APIs are documented according to the OpenAPI standard for information exchange via REST. The entire platform is deployed on the technological infrastructure of the Government of the Balearic Islands, and freely accessible at https://intranet.caib.es/cappescafront/#/Index.

The App was extensively promoted locally through presentations at fishing and yacht clubs, and angler meetings, as well as through radio, print media, and website articles. It also serves as an educational and dissemination tool, providing up-to-date information on policies, fishing seasons, and fishing bans. While the App is freely accessible, the declaration functionalities are only activated after a user has obtained a fishing authorization for an MPA. Self-reporting can be done either by filling in and submitting a printed form or by using the App. Screenshots of the self-reporting process via the App are shown in Figure 2. When using the App, users can declare catches by uploading pictures or by filling in a list of captures (number of individuals per species). In the present study, we considered both the picture and list formats for self-reports, as they were the focus of our analysis. While the list format was automatically organized into spreadsheets, the picture format had to be manually processed into a spreadsheet. A total of 2,562 self-reports were submitted in list format, and 1,110 in picture format.

**Figure 2.**
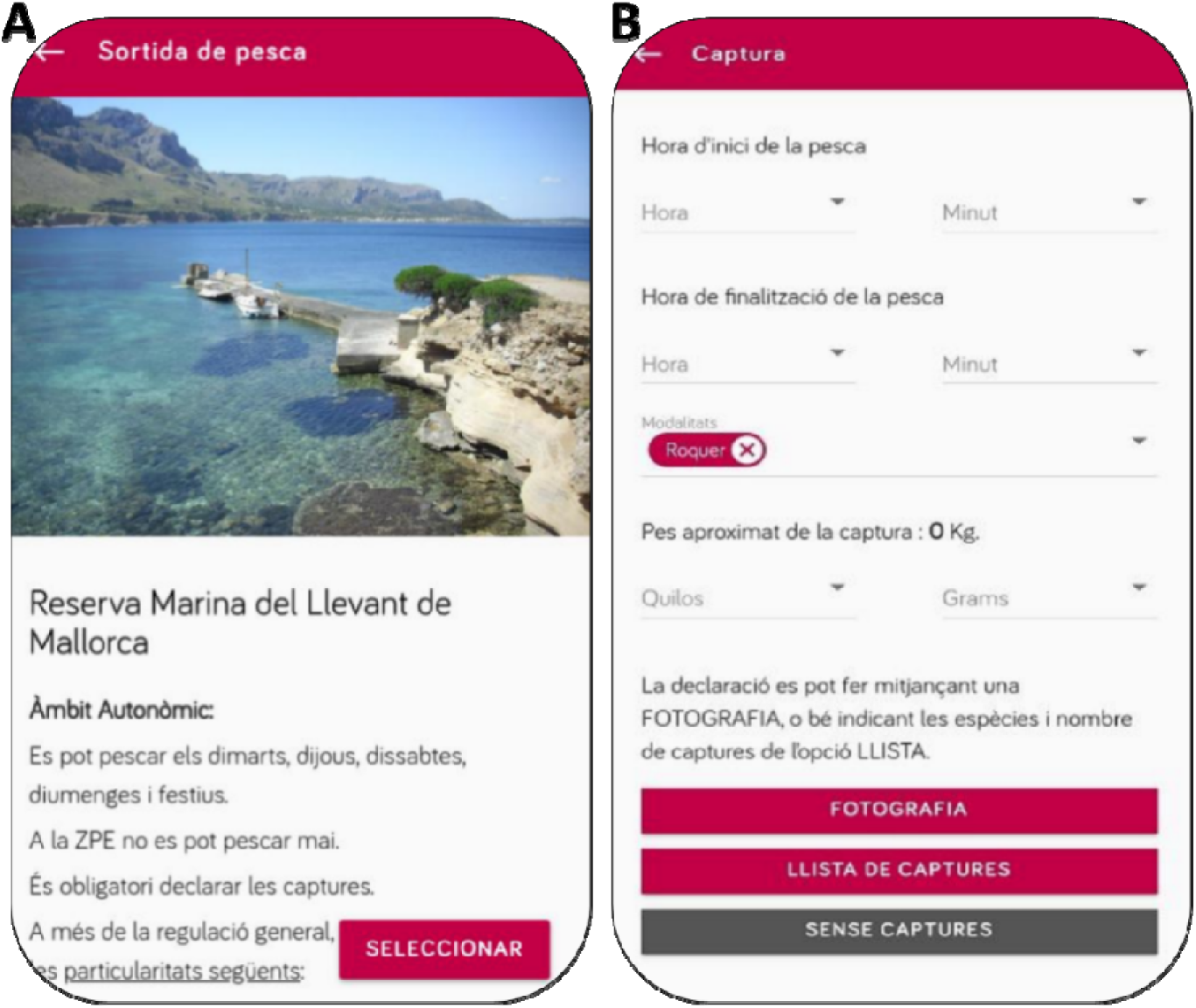
User interface of the angler App “Diari de Pesca Recreativa”. Some self-reporting steps are shown, including (A) Marine Protected Area selection and (B) input of start and finish time, practiced modality, approximate weight of the total catch and link to catch declaration tool (either list or picture).

An App self-report included the following details: the date, start and end times of the fishing trip, the number of anglers on the boat, the name of the MPA, the practiced fishing modality (or modalities), the approximate total weight of the day’s catch, a rough location of the fishing activity, and number of captures for each of a list of 80 species. Self-reports were designed to be completed in under 1 minute, minimizing the burden on anglers. After filtering out unfinished self-reports, with no selected modality or with unrealistic catches, 4,326 self-reports were reduced to 3,672 that were used for analysis, ranging from 08/02/2019 to 15/06/2024 in 12 MPAs and 8 recreational fishing modalities.

### 2.3. On site-creel surveys

In addition to the data collected through the App, a dataset of on-site creel surveys conducted at similar temporal and spatial scales was available for comparison. The creel surveys involved coastguard inspections of recreational fishing boats within MPAs, carried out by MPA coastguards in collaboration with a scientist. To ensure the representation of both seasonal and daily variations in fishing activity, the surveys were distributed across all months, all days of the week, and various times of the day. However, logistical constraints related to inspecting moving boats resulted in a focus on static fishing modalities. Additionally, limitations in work hours led to an overrepresentation of inspections conducted during the morning hours.

During the surveys, anglers were approached respectfully, ensuring minimal disturbance to their fishing activities. In total, 360 boats were inspected over 183 days between 17 July 2017 and 14 May 2023, across 7 MPAs. The creel survey data included the following variables: date, start and inspection times of the fishing trip, duration of the fishing trip up to the point of inspection, the number of anglers on board, the name of the MPA, fishing modality, and captured species along with the number of captures. Sampled MPAs in the Creel survey dataset are limited to “Llevant de Mallorca”, “Sa Dragonera“, “Illes del Toro i Malgrats“, “Costa Nord-Est d’Eivissa-Tagomago“, “Punta de sa Creu“, “Badia de Palma“, “Migjorn de Mallorca“, and “Freus d’Eivissa i Formentera“.

### 2.5. App–Creel survey data comparison and statistical analysis

App and Creel survey datasets were combined to perform data analysis and test differences of E (hour angler · trip) and CPUE (number of captures · E^-1^) estimations at the levels of month, modality, MPA and target species. Variables (i.e. duration, number of anglers, number of captures of a list of 80 species, start and finish trip times) were organized by the App in tabular format.

E is calculated by multiplying the duration of the fishing trip in hours by the number of anglers in each trip:

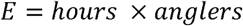

To calculate CPUE, the number of captures is divided by the corresponding trip’s E:

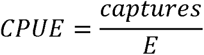

Anglers per hour was calculated from start and finish trip times, which were converted into a matrix of hours (00:00 to 23:00). The corresponding hour was filled with a 1 if trip duration contained that hour and 0 otherwise, which were then multiplied by the number of anglers in that trip.

Generalized Linear Mixed Models (GLMMs) were implemented using the R package glmmTMB (Brooks et al., 2017) to assess gamma-transformed differences in E and CPUE between data sources (App-based self-reports and creel surveys). The models accounted for the non-independence of datasets by treating data from the same MPA as dependent (random effect) when testing the effects of other variables such as month or fishing modality. Analyses were further grouped by month, modality, MPA, and species. Outliers were retained, as all values fell within plausible ranges and were considered valuable for comparing data collection methods. For modality-specific comparisons, where self-reports and inspections recorded multiple fishing modalities per trip, E for each modality was calculated as E = hours x anglers x number of modalities^-1^, while for CPUE = captures x E x number of modalities^-1^. The estimate, standard error and confidence interval of paired groups (App vs. Creel survey) were computed using the function and package emmeans (Lenth et al., 2020) in order to directly statistically test the estimation of E and CPUE. In total, 719 App self-reports and 30 Creel surveys reported more than one practiced modality. R software was used to plot data using R package ggplot2 (Wickham, 2011). A total of 719 App self-reports and 30 creel surveys recorded more than one practiced modality.

## 3. Results

### 3.1. Overview of the App data and performance, and creel survey data

Since the launch of the App, data from 3,672 fishing trips, totalling 11,075 hours of fishing and 54,078 captured fish representing 73 species, were reported by 509 different anglers. Among these self-reports, 575 trips (15.7%) reported no captures. Of the 509 anglers, 175 (34.4%) reported only a single trip. The App reports were distributed across different months, fishing modalities, and marine protected areas (MPAs).

The App accumulated an increasing number of users and self-reports over time (Table 1), following an also increasing number of App users and newly issued MPA fishing authorizations (in August of 2024, 2357 were currently active). The number of self-reporting App users increased each year; however, the growth was outpaced by the increase in the number of potential non-compliant users. This discrepancy led to a significant decline in the self-reporting rate, which dropped to its lowest value of 17.4% in 2023, continuing a downward trend observed since 2021. Potential non-compliant users fall into two indistinguishable categories in the dataset: false non-compliants, anglers who received authorization to fish but never actually fished in the MPA and, therefore, did not contribute to fishing effort (E); and true non-compliants, anglers who failed to report their fishing trips in MPAs, either knowingly or unknowingly, resulting in unaccounted fishing effort. It is important to note that the number of authorizations and active users does not align because individual users can hold authorizations for multiple MPAs.

**Table 1.**
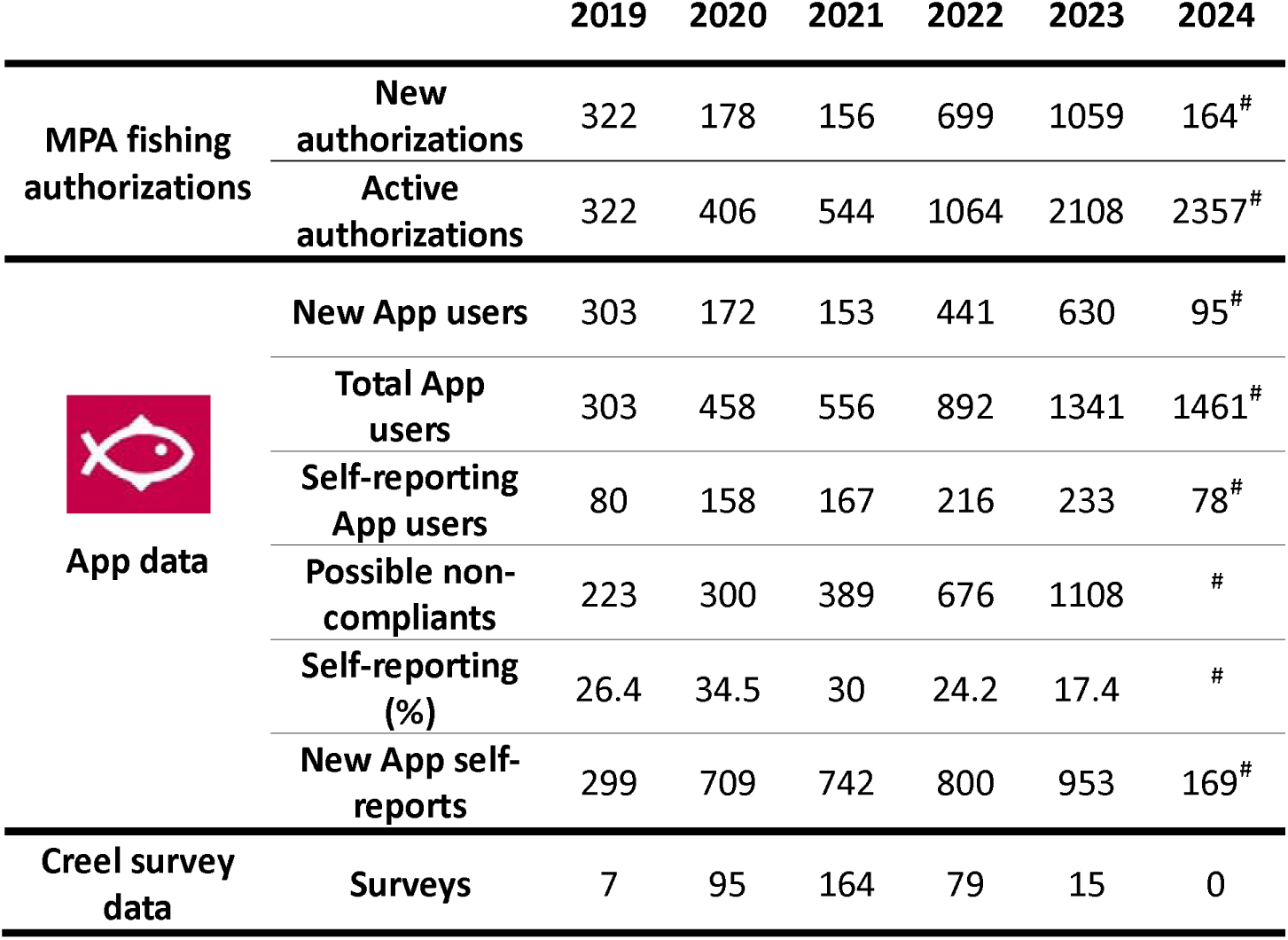
App users, App declaration data and Creel survey inspections over time. ^#^Year 2024 is not finished.

|  |  | 2019 | 2020 | 2021 | 2022 | 2023 | 2024 |
| --- | --- | --- | --- | --- | --- | --- | --- |
| MPA fishing<br>authorizations | New<br>authorizations | 322 | 178 | 156 | 699 | 1059 | 164<sup>#</sup> |
|  | Active<br>authorizations | 322 | 406 | 544 | 1064 | 2108 | 2357<sup>#</sup> |
| App data | New App users | 303 | 172 | 153 | 441 | 630 | 95<sup>#</sup> |
|  | Total App<br>users | 303 | 458 | 556 | 892 | 1341 | 1461<sup>#</sup> |
|  | Self-reporting<br>App users | 80 | 158 | 167 | 216 | 233 | 78<sup>#</sup> |
|  | Possible non-compliance | 223 | 300 | 389 | 676 | 1108 | # |
|  | Self-reporting<br>(%) | 26.4 | 34.5 | 30 | 24.2 | 17.4 | # |
|  | New App self-reports | 299 | 709 | 742 | 800 | 953 | 169<sup>#</sup> |
| Creel survey<br>data | Surveys | 7 | 95 | 164 | 79 | 15 | 0 |

Creel survey E data from 360 fishing trips, 673 hours of fishing and 4176 captured fish from 59 species was obtained. Of these inspections, 22 had 0 captures (6.11%). Creel surveys were distributed across months, fishing modalities and MPAs.

### 3.2. Estimates of Recreational fishing Effort (E) per month, modality, and MPA

Figure 3 offers a view of the raw data. It shows a clear trend of higher overall values of E in App data (4.74 ± 3.9 SD hours · angler · trip) than in Creel Survey data (3.20 ± 2.4 SD hours · angler · trip), which presented significant differences according to GLMM (p < 0.001) (Figure 4, Table 2). When analysing the most extreme outliers, no clear association with specific months or MPAs was observed. Instead, the modalities of squid fishing, trolling fishing, and shallow fishing accounted for the majority of these outliers (Figure 3).

**Figure 3.**
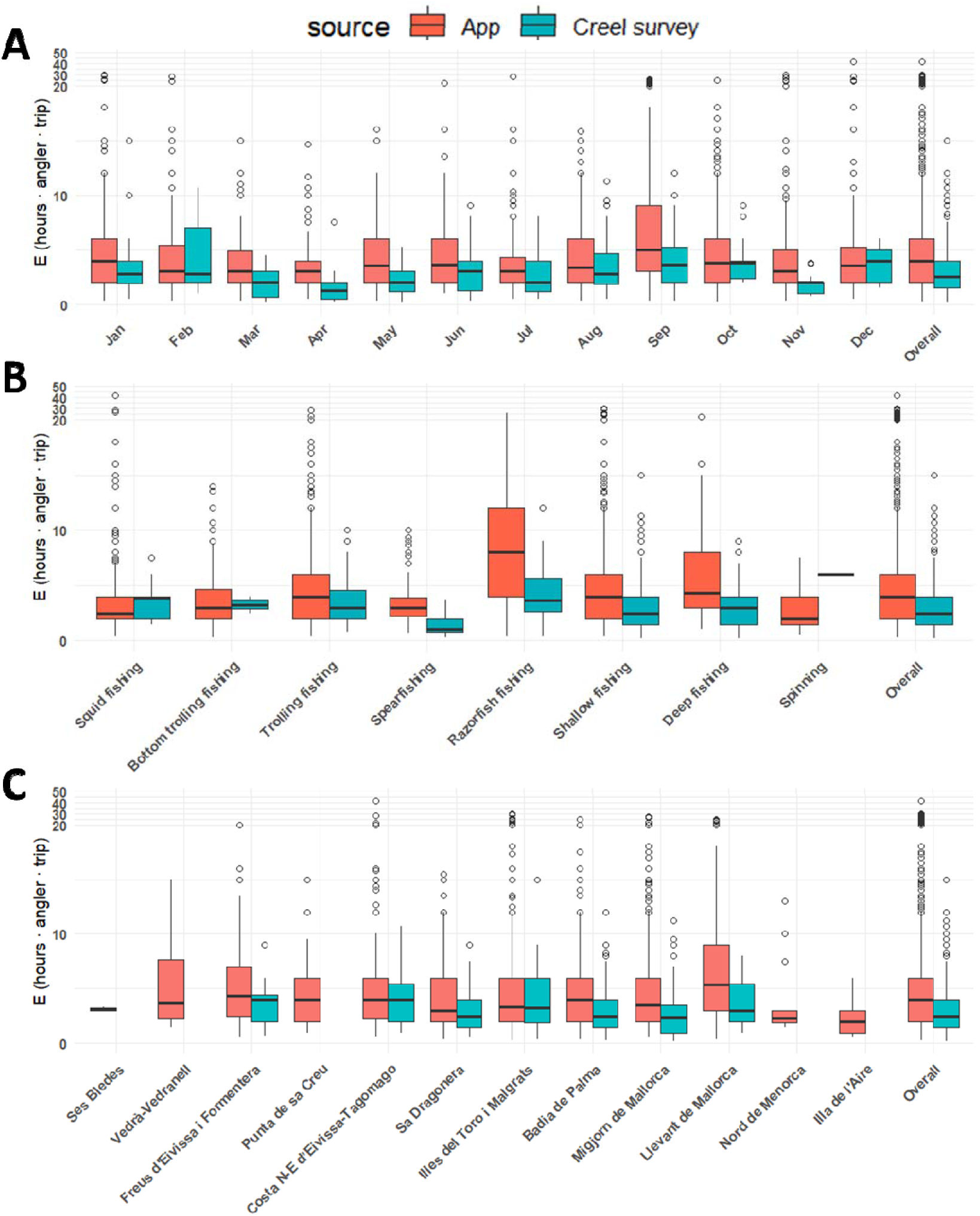
Effort in App data and Creel survey data for each month (A), fishing modality (B) and Balearic marine protected area (C). Overall group represents whole datasets without grouping, and is shown in all panels for reference. Statistic test results are shown in table 2. The y-axis above 20 is divided by 10 for visualization purposes. Boxplots indicate upper limit and lower limit of data excluding outliers (whiskers), third quartile (top of box), median (thick line) and first quartile (bottom of box).

**Figure 4.**
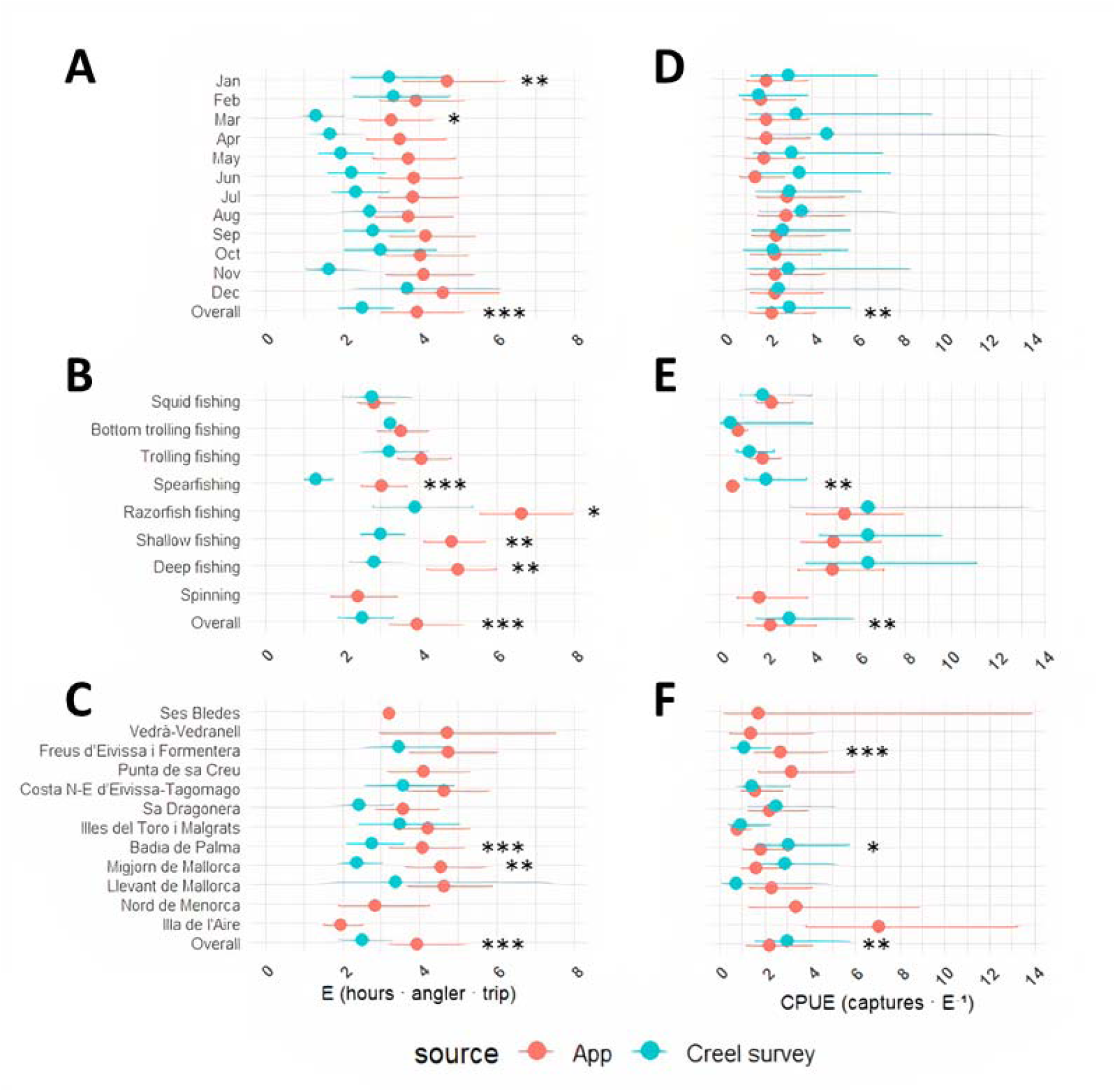
Generalized Linear Mixed Model results of Effort (E, left) and Captures per Unit of Effort (CPUE, right) in App data and Creel survey data for each month (A & D), fishing modality (B & E) and Balearic marine protected area (C & F). Overall group represents whole datasets without grouping, and is shown in all panels for reference. Statistic results are shown in table 2. Points indicate the mean of the response and whiskers show upper and lower confidence limits.

**Table 2.** Generalized Linear Mixed Model results for the statistical comparison of Effort (hours · angler · trip) estimations between App data and Creel survey data without grouping (overall) and by month, modality and marine protected area (MPA). App n shows the number of self-reports used in the calculation and Creel survey n the number of surveys. SE: Standard error. Significance codes for p-value (< 0.001: ***, < 0.01: **, < 0.05: *) are shown in significantly different comparisons.

|  | <u>App</u> |  |  | <u>Creel survey</u> |  |  | <u>Comparison</u> |  |
| --- | --- | --- | --- | --- | --- | --- | --- | --- |
|  | n | Estimate | SE | n | Estimate | SE | Z-value | p-value |
| **Overall E** | *Model: CPUE ~ source \* (1 \| MPA) + (1 \| modality) + (1 \| month)* |  |  |  |  |  |  |  |
| Overall | 3672 | 3.92 | 0.5 | 360 | 2.50 | 0.4 | -11.62 | <0.001*** |
| **by month** | *CPUE ~ source \* month + (1 \| modality) + (1 \| MPA)* |  |  |  |  |  |  |  |
| January | 272 | 4.70 | 0.7 | 28 | 3.20 | 0.6 | -2.95 | 0.003** |
| February | 239 | 3.90 | 0.5 | 25 | 3.31 | 0.6 | 1.15 | 0.251 |
| March | 146 | 3.26 | 0.5 | 13 | 1.32 | 0.3 | -2.25 | 0.024* |
| April | 95 | 3.49 | 0.5 | 17 | 1.69 | 0.4 | -1.57 | 0.116 |
| May | 115 | 3.70 | 0.5 | 30 | 1.96 | 0.4 | -1.34 | 0.180 |
| June | 170 | 3.85 | 0.5 | 45 | 2.24 | 0.4 | -0.92 | 0.360 |
| July | 371 | 3.83 | 0.5 | 62 | 2.34 | 0.4 | -0.67 | 0.502 |
| August | 461 | 3.71 | 0.5 | 44 | 2.70 | 0.5 | 0.41 | 0.680 |
| September | 767 | 4.17 | 0.6 | 54 | 2.80 | 0.5 | -0.09 | 0.932 |
| October | 499 | 4.02 | 0.5 | 20 | 3.00 | 0.6 | 0.45 | 0.651 |
| November | 216 | 4.11 | 0.6 | 13 | 1.64 | 0.4 | -2.34 | 0.019* |
| December | 319 | 4.61 | 0.6 | 9 | 3.68 | 0.9 | 0.61 | 0.543 |
| **by modality** | *Model: CPUE ~ source \* modality + (1 \| month) + (1 \| MPA)* |  |  |  |  |  |  |  |
| Squid | 1016 | 2.83 | 0.2 | 22 | 2.75 | 0.5 | -0.19 | 0.849 |
| Bottom trolling fishing | 373 | 3.51 | 0.3 | 3 | 3.24 | 1.5 | -0.11 | 0.915 |
| Trolling fishing | 828 | 4.05 | 0.4 | 38 | 3.21 | 0.4 | -1.14 | 0.254 |
| Spearfishing | 167 | 3.02 | 0.3 | 36 | 1.31 | 0.2 | -4.3 | <0.001*** |
| Razorfish fishing | 516 | 6.65 | 0.6 | 22 | 3.87 | 0.6 | -2.52 | 0.012 * |
| Shallow fishing | 1187 | 4.83 | 0.4 | 194 | 2.99 | 0.3 | -2.96 | 0.003 ** |
| Deep fishing | 555 | 5 | 0.5 | 81 | 2.82 | 0.4 | -3.12 | 0.002 ** |
| Spinning | 29 | 2.41 | 0.4 | 1 | NA | NA | NA | NA |
| **by MPA** | *Model: CPUE ~ source \* MPA + (1 \| modality) + (1 \| month)* |  |  |  |  |  |  |  |
| Ses Bledes | 2 | 3.22 | 1.5 | 0 | NA | NA | NA | NA |
| Vedrà-Vedranell | 10 | 4.72 | 1.1 | 0 | NA | NA | NA | NA |
| Freus d'Eivissa i Formentera | 257 | 4.74 | 0.6 | 37 | 3.45 | 0.5 | 0.49 | 0.628 |
| Punta de sa Creu | 96 | 4.10 | 0.5 | 0 | NA | NA | NA | NA |
| C. N-E d'Eivissa-Tagomago | 485 | 4.63 | 0.5 | 33 | 3.56 | 0.6 | 0.86 | 0.393 |
| Sa Dragonera | 336 | 3.58 | 0.4 | 40 | 2.43 | 0.4 | 0.14 | 0.988 |
| Illes del Toro i Malgrats | 473 | 4.22 | 0.5 | 20 | 3.49 | 0.7 | 1.15 | 0.252 |
| Badia de Palma | 252 | 4.06 | 0.5 | 83 | 2.75 | 0.4 | -4.64 | <0.001*** |
| Migjorn de Mallorca | 1339 | 4.56 | 0.5 | 144 | 2.37 | 0.3 | -2.61 | 0.009 ** |
| Llevant de Mallorca | 302 | 4.64 | 0.6 | 3 | 3.39 | 1.3 | 0.18 | 0.854 |
| Nord de Menorca | 15 | 2.85 | 0.6 | 0 | NA | NA | NA | NA |
| Illa de l'Aire | 105 | 1.95 | 0.3 | 0 | NA | NA | NA | NA |

Grouping of the data showed that summer months (July, August, September) and winter (December, January) presented the longest durations and/or number of anglers per trip for both data sources (Figure 3A). The highest values were recorded in September. Accumulating the E of 767 self-reports for App data and 54 inspections for Creel survey, September recorded a summatory of E of 4821 hours · angler · trip for App data and 215 for Creel survey data, followed by October, August and July. The lowest values were recorded in April, with 357 and 32 hours · angler · trip, respectively. GLMM results showed significant differences between App data and Creel survey data were present in January, March and November, in all cases presenting smaller E estimates by the creel survey (Figure 4A, Table 2).

Estimation of average E across fishing modalities (Figure 3B) showed that Razorfish fishing had the highest average values of E for both datasets. As summatory of E, it accumulated 3401 hours · angler · trip for App data and 96.3 for Creel survey data. This is still lower than the summatory of E values for Squid fishing, with E of 3431 and 76.2, respectively, and Shallow fishing with 4369 and 618, respectively. Trolling fishing was also worthy of mention, with 3244 and 130 hours · angler · trip, respectively, and Deep fishing with 1606 and 165, respectively. For Trolling fishing, 68.05% of E was concentrated in September and October. App data presented significantly higher E estimations in modalities Spearfishing, Razorfish, Shallow, and Deep fishing (Figure 4B, Table 2). It is relevant to point out that for Spearfishing, only 20 App users declared and one of them accumulated 45.37% of the E. This same user accumulated 29.03% of the Spearfishing captures. It is also worth noting that Shallow and Deep fishing are the two static modalities that are practiced all year round and they are conformed of 194 and 81 inspections, representing 76.39% of the Creel survey inspections. Logistic problems with the surveying of other modalities lead to the decision to focus on these two.

In general, durations and/or number of anglers per trip were similar among MPAs (Figure 3C), although some MPAs received more fishing pressure than others: “Migjorn de Mallorca” (1339 self-reports), “Illes del Toro i Malgrats” (473 self-reports), and “Costa Nord-Est d’Eivissa-Tagomago” (485 self-reports) received the most fishing activity. “Ses Bledes” (2 self-reports), “Vedrà-Vedranell” (10 self-reports), and “Nord de Menorca” (15 self-reports) received the least. When estimating E, significant differences were detected only in the MPAs “Migjorn de Mallorca” and “Badia de Palma”, with higher values for App data (Figure 4C, Table 2). Importantly, “Migjorn de Mallorca” MPA has the highest number of App self-reports (1339, 36.47% of the total) and of Creel surveys (144, 40% of the total), which translates into a total E of 6073 and 251.57 hours · angler · trip, respectively. Due to the high portion of the data it contains, the difference in this specific MPA has a large input in the differences in the overall datasets comparison (Table 2).

The format in which both data sources were collected allowed for the visualization of E at different times of the day. In Figure 5A (App data) and 5B (Creel survey data), the summatory of E per hour of the day over the entire period covered by the datasets (App: 2019-2024, Creel survey: 2019 - 2023) is displayed, showing that the most fishing activity is usually performed between 9:00 am and 12:00 pm for App users and slightly more spread for Creel survey. Fishing modalities have peak activity times at different times of the day. The case of Squid fishing is especially interesting, with peak activities in the morning and afternoon, aiming for the most active hours of the target species. As mentioned above, Shallow, Squid, Razorfish and Trolling fishing modalities accumulated the most summatory of E. It is important to note that only some Creel survey inspections collected time at start of the fishing trip (Spinning: 1, Deep: 2, Shallow: 76, Razorfish: 4, Spearfishing: 36, Trolling: 24, Bottom trolling: 3, Squid: 22) and therefore some modalities are not well-represented.

**Figure 5.**
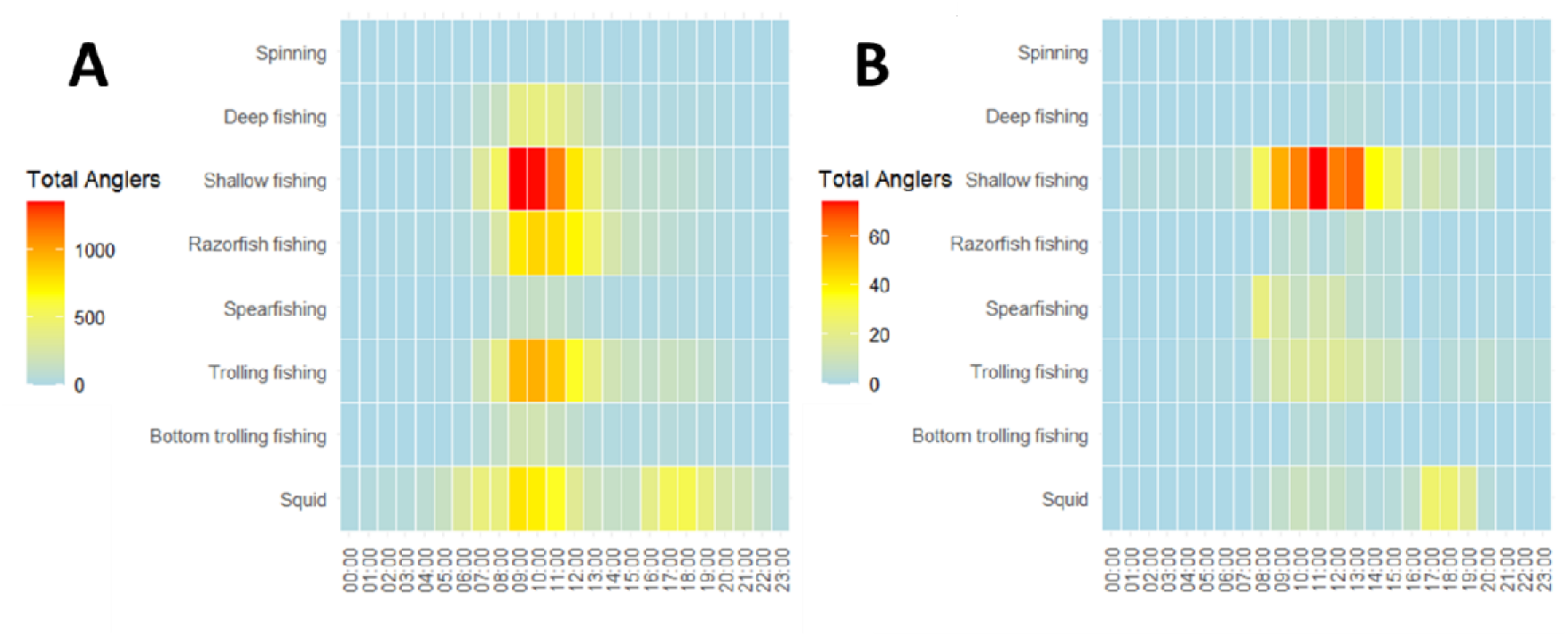
Summatory of anglers per fishing modality in each hour of the day over the entire period covered by each dataset: A) App data and B) Creel survey data. The legend shows the intensity of summatory of anglers per hour.

### 3.3. Estimates of Catch per Unit of Effort (CPUE) per month, modality, MPA, and species

Figure 6 shows CPUE data prior to GLMM treatment. App data (2.95 ± 3.9 SD captures · E^-1^) present lower overall values of CPUE than Creel Survey data (4.23 ± 4.6 SD captures · E^-1^), which is confirmed by significant differences in GLMM testing (p = 0.001) (Figure 4, Table 3). Similar to E, extreme outliers were more commonly associated to modalities Trolling, Razorfish, Shallow and Deep fishing than to specific months or MPAs.

**Figure 6.**
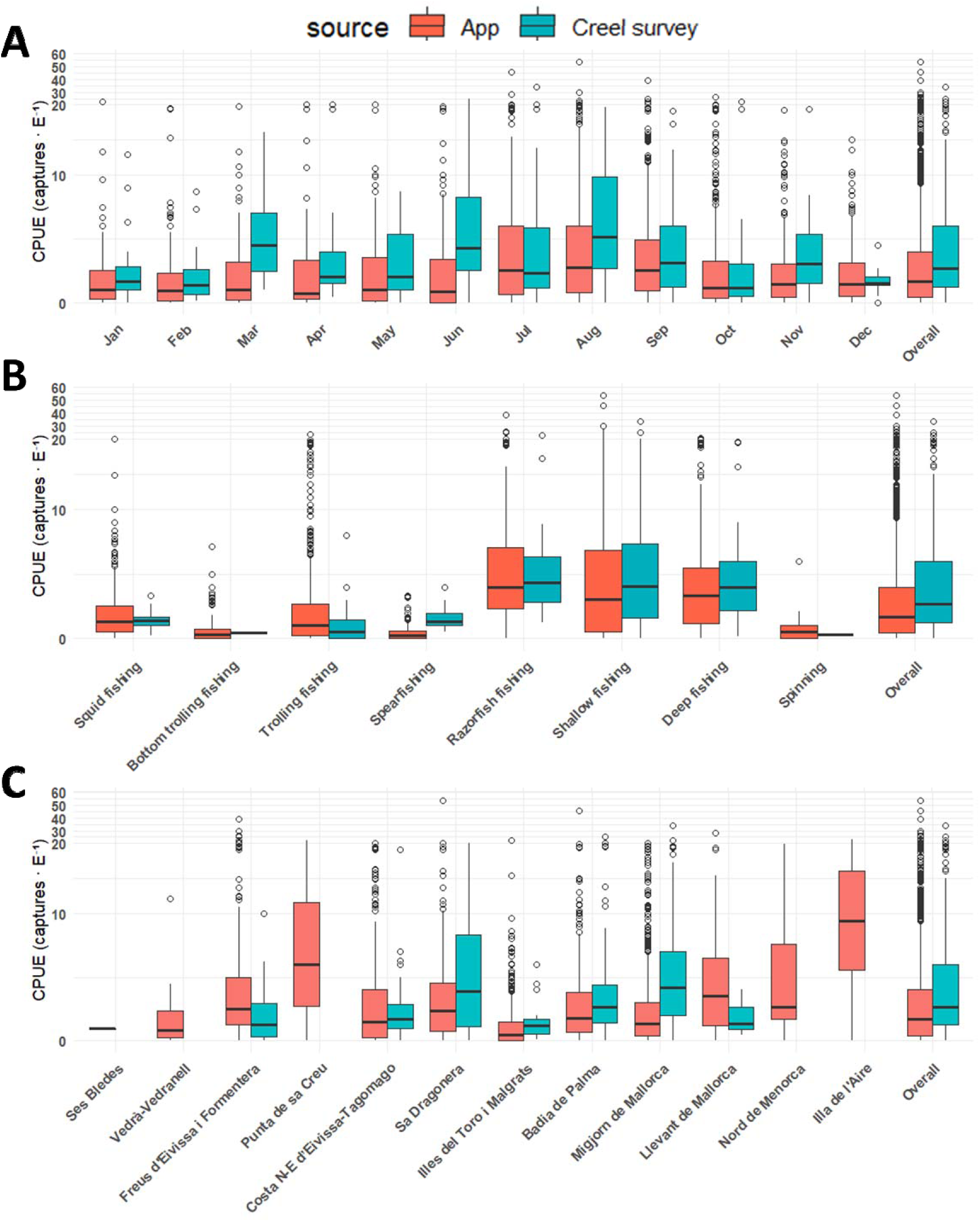
Catch per Unit of Effort (CPUE) in App data and Creel survey data for each month (A), fishing modality (B) and Balearic marine protected area (C). Overall group represents whole datasets without grouping, and is shown in all panels for reference. Statistic test results are shown in table 3. The y-axis above 20 is divided by 10 for visualization purposes. Boxplots indicate upper limit and lower limit of data excluding outliers (whiskers), third quartile (top of box), median (thick line) and first quartile (bottom of box).

**Table 3.** Generalized Linear Mixed Model results for the statistical comparison of Captures per Unit of Effort (CPUE) estimations in Captures · E between App data and Creel survey data without grouping (overall) and by month, modality and marine protected area (MPA). App n shows the number of self-reports used in the calculation and Creel survey n the number of surveys. SE: Standard error. Significance codes for p-value (< 0.001: ***, < 0.01: **, < 0.05: *) are shown in significantly different comparisons.

|  | App |  |  | Creel survey |  |  | Comparison |  |
| --- | --- | --- | --- | --- | --- | --- | --- | --- |
|  | n | Estimate | SE | n | Estimate | SE | Z-value | p-value |
| **Overall CPUE** | *Model: CPUE ~ source \* (1 \| MPA) + (1 \| modality) + (1 \| month)* |  |  |  |  |  |  |  |
| Overall | 3672 | 2.20 | 0.7 | 360 | 2.99 | 1.00 | 3.20 | 0.001** |
| **by month** | *Model: CPUE ~ source \* month + (1 \| modality) + (1 \| MPA)* |  |  |  |  |  |  |  |
| January | 272 | 1.95 | 0.7 | 28 | 2.94 | 1.3 | 1.29 | 0.196 |
| February | 239 | 1.66 | 0.6 | 25 | 1.58 | 0.7 | -1.02 | 0.310 |
| March | 146 | 1.95 | 0.7 | 13 | 3.27 | 1.8 | 0.20 | 0.839 |
| April | 95 | 1.92 | 0.7 | 17 | 4.70 | 2.3 | 0.94 | 0.346 |
| May | 115 | 1.82 | 0.6 | 30 | 3.08 | 1.3 | 0.26 | 0.792 |
| June | 170 | 1.41 | 0.5 | 45 | 3.45 | 1.4 | 1.19 | 0.233 |
| July | 371 | 2.86 | 0.9 | 62 | 2.97 | 1.1 | -0.97 | 0.331 |
| August | 461 | 2.85 | 0.9 | 44 | 3.52 | 1.4 | -0.49 | 0.621 |
| September | 767 | 2.40 | 0.8 | 54 | 2.69 | 1.0 | -0.76 | 0.446 |
| October | 499 | 2.32 | 0.8 | 20 | 2.21 | 1.1 | -0.96 | 0.336 |
| November | 216 | 2.34 | 0.8 | 13 | 2.94 | 1.1 | -0.33 | 0.739 |
| December | 319 | 2.33 | 0.8 | 9 | 2.46 | 1.5 | -0.58 | 0.565 |
| **by modality** | *Model: CPUE ~ source \* modality + (1 \| month) + (1 \| MPA)* |  |  |  |  |  |  |  |
| Squid | 1016 | 2.21 | 0.4 | 22 | 1.87 | 0.7 | -0.49 | 0.624 |
| Bottom trolling fishing | 373 | 0.8 | 0.2 | 3 | 0.44 | 0.5 | -0.36 | 0.717 |
| Trolling fishing | 828 | 1.84 | 0.3 | 38 | 1.26 | 0.4 | -0.5 | 0.62 |
| Spearfishing | 167 | 0.56 | 0.1 | 36 | 2.01 | 0.6 | 3.26 | 0.001** |
| Razorfish fishing | 516 | 5.41 | 1 | 22 | 6.42 | 2.4 | 0.7 | 0.484 |
| Shallow fishing | 1187 | 4.91 | 0.9 | 194 | 6.42 | 1.3 | 1.2 | 0.232 |
| Deep fishing | 555 | 4.89 | 0.9 | 81 | 6.42 | 1.8 | 1.05 | 0.293 |
| Spinning | 29 | 1.7 | 0.7 | 1 | NA | NA | NA | NA |
| **by MPA** | *Model: CPUE ~ source \* MPA + (1 \| modality) + (1 \| month)* |  |  |  |  |  |  |  |
| Ses Bledes | 2 | 1.71 | 2.0 | 0 | NA | NA | NA | NA |
| Vedrà-Vedranell | 10 | 1.36 | 0.8 | 0 | NA | NA | NA | NA |
| Freus d'Eivissa i Formentera | 257 | 2.69 | 0.8 | 37 | 1.07 | 0.4 | -4.20 | <0.001*** |
| Punta de sa Creu | 96 | 3.18 | 1.0 | 0 | NA | NA | NA | NA |
| C. N-E d'Eivissa-Tagomago | 485 | 1.59 | 0.5 | 33 | 1.44 | 0.6 | -1.78 | 0.076 |
| Sa Dragonera | 336 | 2.23 | 0.6 | 40 | 2.50 | 0.9 | -1.21 | 0.227 |
| Illes del Toro i Malgrats | 473 | 0.81 | 0.2 | 20 | 0.93 | 0.4 | -0.91 | 0.362 |
| Badia de Palma | 252 | 1.82 | 0.5 | 83 | 3.05 | 1.0 | 2.53 | 0.011* |
| Migjorn de Mallorca | 1339 | 1.62 | 0.5 | 144 | 2.88 | 0.9 | 0.25 | 0.803 |
| Llevant de Mallorca | 302 | 2.33 | 0.7 | 3 | 0.76 | 0.7 | -1.76 | 0.079 |
| Nord de Menorca | 15 | 3.39 | 1.7 | 0 | NA | NA | NA | NA |
| Illa de l'Aire | 105 | 7.10 | 2.3 | 0 | NA | NA | NA | NA |

Opposite to E, CPUE across months was generally slightly higher in Creel survey data (Figure 6A) but with no significant differences (Figure 4D, Table 3). The highest values were recorded in summer for both datasets. By accumulation of captures, the top month was September with 19520 fish caught by App users and 762 in Creel survey data. August saw 8264 and 849 fish caught, respectively, and October 6840 and 208, respectively. The lowest month was April, with 924 and 98 fish caught, respectively.

Estimation of CPUE across fishing modalities (Figure 6B) showed that Razorfish fishing and Shallow fishing have the highest catch rate per trip, followed by Deep, Trolling and Squid fishing. The accumulated number of captures of these modalities in App data was respectively 16411, 17525, 5640, 7177, and 6526. For Creel survey data they were respectively 466, 2754, 636, 134, and 105. Significant differences among data sources were only present in the modality Spearfishing (p = 0.001) (Figure 4E, Table 3), with higher values in Creel survey.

Comparison across MPAs (Figure 6C) showed that some of the least exploited MPAs presented the highest CPUE: “Nord de Menorca”, “Punta de sa Creu” and “Illa de l’Aire”. Significant differences were detected in only two MPAs: For “Freus d’Eivissa i Formentera” App data was highest, while for “Migjorn de Mallorca” Creel survey data was highest (Figure 4F, Table 3).

CPUE was also calculated for 8 species of especial interest (Figure 7). CPUE was normally higher in Creel survey data except for *Loligo vulgaris* but significant differences among data sources were detected in only half of the species: *Coris julis*, *Diplodus vulgaris*, *Loligo vulgaris*, and *Serranus scriba* (Table 4). Differences in mean CPUE values per species were higher in Creel survey data by 1.7 individuals per fishing trip for *Coris julis*, 1.6 for *Diplodus annularis*, 1.6 for *Diplodus vulgaris*, 1 for *Seriola dumerili*, 0.8 for *Serranus cabrilla*, 1.7 for *Serranus scriba*, and 0.3 for *Xyrichthys novacula*. L*oligo vulgaris* was 0.2 individuals per fishing trip higher in App data. Differences in median CPUE values per species were higher in Creel survey data by 1.5 individuals per fishing trip for *Coris julis*, 1.1 for *Diplodus annularis*, 1 for *Diplodus vulgaris*, 0 for *Loligo vulgaris*, 0.4 for *Seriola dumerili*, 1.5 for *Serranus cabrilla*, 2.5 for *Serranus scriba*, and 0.8 for *Xyrichthys novacula. Loligo vulgaris* was among the top species in number of caught individuals (App: 5193, Creel survey: 110). *Xyrichthys novacula* was the top captured species (App: 13820, Creel survey: 339). Of these *X. novacula*, 11364 (App) and 251 (Creel survey) were captured in September. *Serranus scriba* (App: 10415, Creel survey: 1090) was the top Shallow fishing species, while *Diplodus annularis*(App: 1892, Creel survey: 388) and *Coris julis* (App: 1457, Creel survey: 314) are also very common captures of the modality. *Seriola dumerili* (App: 1089, Creel survey: 37) is the top Trolling fishing species. *Serranus cabrilla* is the target species and top captured species of Deep fishing (App: 10156, Creel survey: 904). *Diplodus vulgaris* (App: 684, Creel survey: 126) is a relatively common capture of both Deep and Shallow fishing.

**Figure 7.**
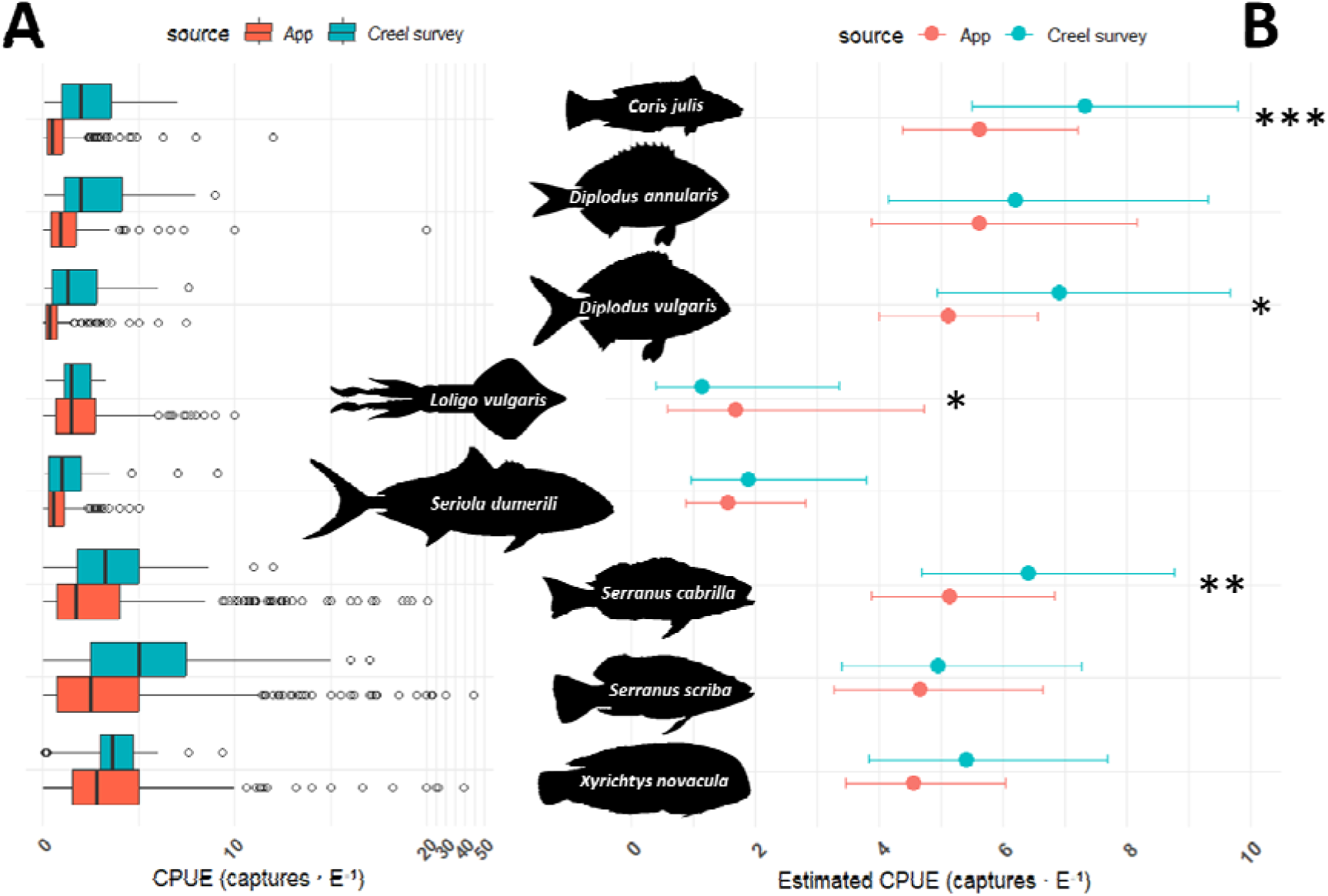
Catch per Unit of Effort (CPUE) for 8 species of interest calculated from App data and Creel survey data. (A) Untreated data comparison. The y-axis above 20 is divided by 10 for visualization purposes. Boxplots indicate upper limit and lower limit of data excluding outliers (whiskers), third quartile (top of box), median (thick line) and first quartile (bottom of box). (B) Generalized Linear Mixed Model results. Points indicate the mean of the response and whiskers show upper and lower confidence limits. Statistic test results are shown in table 4.

**Table 4.** Generalized Linear Mixed Model results for the statistical comparison of Catch per Unit of Effort (CPUE) between App data and Creel survey data for 8 species of interest. The columns n shows the number of self-report/creel surveys used in the calculation while the columns titled fish show the number of captures for each species and data source. SE: Standard error. Significance codes for p-value (< 0.001: ***, < 0.01: **, < 0.05: *) are shown in significantly different comparisons.

| Species | App |  |  |  | Creel survey |  |  |  | Comparison |  |
| --- | --- | --- | --- | --- | --- | --- | --- | --- | --- | --- |
|  | n | Fish | Estimate | SE | n | Fish | Estimate | SE | Z value | p-value |
|  | *CPUE ~ source + (1 \| MPA) + (1 \| modality) + (1 \| month)* |  |  |  |  |  |  |  |  |  |
| *Coris julis* | 314 | 1457 | 5.62 | 0.7 | 84 | 330 | 7.33 | 1.1 | 3.30 | <0.001*** |
| *Diplodus annularis* | 388 | 1892 | 5.63 | 1.1 | 89 | 348 | 6.21 | 1.3 | 1.18 | 0.236 |
| *Diplodus vulgaris* | 126 | 684 | 5.12 | 0.6 | 45 | 196 | 6.91 | 1.2 | 2.27 | 0.023* |
| *Loligo vulgaris* | 110 | 5193 | 1.68 | 0.9 | 24 | 869 | 1.14 | 0.6 | -2.39 | 0.017* |
| *Seriola dumerili* | 37 | 1089 | 1.57 | 0.5 | 19 | 266 | 1.9 | 0.7 | 0.90 | 0.370 |
| *Serranus cabrilla* | 904 | 10156 | 5.15 | 0.7 | 131 | 725 | 6.41 | 1.0 | 3.05 | 0.002** |
| *Serranus scriba* | 1090 | 10415 | 4.66 | 0.8 | 136 | 637 | 4.96 | 1.0 | 0.88 | 0.378 |
| *Xyrichthys novacula* | 339 | 13820 | 4.57 | 0.6 | 32 | 489 | 5.42 | 1.0 | 1.37 | 0.171 |

## Discussion

The interest in digital platforms to obtain data for recreational fisheries science is growing, but the intrinsic avidity bias in the data and general concerns on the validity of self-reported citizen science data tend to prevent the meaningful use of this data for scientific research or policy making (Lennox et al., 2022). Here we provide an analysis of the performance of the App in reporting recreational fisheries data over a six-years period in the Balearic Island. We have found that self-reported data from the app demonstrated significant differences in fishing effort (E) and captures per Unit of Effort (CPUE) estimates compared to traditional creel surveys, with the App tending to overestimate effort and creel surveys underestimating it. Despite these differences, most comparisons across species, fishing modalities, months, and Marine Protected Areas (MPAs) showed no statistically significant variation in E and CPUE estimates. Integrating data from both methods enhances the accuracy and reliability of recreational fisheries monitoring, leveraging the app’s broader data collection capacity and automation potential.

Non-digital, paper-based self-reporting of catches via fishing diaries has been widely evaluated as a source of recreational fisheries data (Bellanger & Levrel, 2017; Rocklin et al., 2014), but the newer digital formats have received less attention (Griffiths et al., 2013; Skov et al., 2021; Venturelli et al., 2017), with few exceptions (Jiorle et al., 2016; Johnston et al., 2022; Papenfuss et al., 2015). Previous work has been published on the validity of data obtained from recreational angler online surveys and apps; some used logical reasoning (Skov et al., 2021; Venturelli et al., 2017) and others experimentally tested it (Griffiths et al., 2013; Jiorle et al., 2016; Johnston et al., 2022; Papenfuss et al., 2015). When recreational anglers self-report their fishing success out of voluntariness to collaborate, data is biased towards an overrepresentation of avid personality types, leading to biased data (Griffiths et al., 2013; Gundelund et al., 2020; Trinnie & Ryan, 2024). Empirically, after testing time-location sampling, access-point survey and online diaries for the surveying of recreational longtail tuna (*Thunnus tonggol*) catches, the online method was dismissed due to avidity, volunteerism, and differential recruitment bias (Griffiths et al., 2013). Other studies would have a positive conclusion on the use of app data provided bias-mitigation measures were implemented (Gundelund et al., 2020; Trinnie & Ryan, 2024). Studies with a positive outcome include Johnston et al. (2022), who showed that data from the app MyCatch, after careful curation and for specific species and regions, could provide catch rate estimates comparable to those obtained from conventional fisheries-dependent surveys (i.e. national mailing survey, creel surveys and gillnetting surveys) but was not suitable to estimate fish abundance or relative composition of caught species. The app MyCatch is aimed at anglers who want to be alerted about events like fishing competitions or want to collaborate with researchers. Papenfuss et al. (2015) were able to observe angler preferences and movement networks despite obtaining data from an app (Ifish Alberta) which was not developed for research. Jiorle et al. (2016) showed that median catch rate for particular species could be comparable between app (iAngler) data and conventional sources (Marine Recreational Information Program survey of Florida).

App data records start in 2019, with the incorporation of “Llevant de Mallorca” MPA into the declaration system. Other MPAs were progressively incorporated over the course of 2019-2023 until all 12 Balearic MPAs were present in the App, although some of them still have few self-reports. The number of issued authorizations, App users and App self-reports increased steadily over the years, reaching a total of 2357, 1164 and 4326, respectively, in August of 2024. The number of possible non-declarants also increased reaching 1108 in 2023, or 82.62% of total App users; this means that early users from 2019-2021 were more likely to declare that subsequently incorporated users. These users do not declare because they do not go fishing in MPAs despite having the authorization, out of non-conformity or because they do not know about the App and current regulations. Strategies such as incentives, user engagement enhancements, and dissemination work can help improve participation (Gundelund et al., 2020) and will be carried out for the App. Presentations in fishing clubs and angler associations are especially important to reach the biggest number of anglers. Regardless, the actual number of anglers who go fishing and do not declare - which do represent a problem for both E and CPUE estimation - is unknown but likely much smaller than the worst-case scenario that 82.62% represents.

Cornesse et al. (2020) emphasized the need for studies that compare opt-in app data with data obtained from traditional surveys in order to assess the reliability of E and CPUE. In our study, differences between App and Creel survey data were compared. E was overall significantly higher in App data while CPUE was significantly higher in Creel survey data. The sources of these differences were probably related to the limitations of each data collection method. For Apps in general, (1) When obtaining App data for species subject to a season ban or limited to one specimen per angler and day (e.g. *Scorpaena scrofa, Dentex dentex, Sciaena umbra, Sparus aurata, Zeus faber, Epinephelus marginatus*), scepticism is advised. For this our specific App, (2) some users may have declared their start time as the hour when they turn on their boat, not when they start fishing. And likewise with the finish hour, thus declaring an unrealistic fishing trip time and estimate of E. t is not possible to determine how many users followed proper self-reporting practices; however, future app dissemination efforts will emphasize correct usage, and app functionalities will be revised to improve clarity and user compliance. Regarding the limitations of creel surveys, (1) the sampling methodology was constrained by work-time schedules, resulting in overrepresentation of some time periods while others were underrepresented, and (2) a relatively limited number of surveys were conducted due to their (3) time-intensive nature and (4) high cost (Lewin et al., 2021). Additionally, (5) creel surveys, conducted during ongoing fishing trips rather than at their conclusion, tended to underestimate trip durations. Finally, (6) creel surveys exhibited modality bias, as fishing modalities practiced statically and year-round (e.g., shallow and deep fishing) were easier to survey and accounted for 76.39% of the inspections. Both data collection methods were designed to minimize disturbance to anglers, limiting the duration of surveys and information collected. While creel surveys required 5–10 minutes and temporarily interrupted anglers’ activities, experienced app users could complete their declarations in under one minute, reducing time investment.

Regarding monthly estimates, both average and summatory of E were highest in September for both datasets. This was in part due to the lifting of the Razorfish fishing ban on the 1^st^ of September. The fishing season of other species also explained high average and summatory of E in other months, like Squid fishing during winter. Monthly CPUE is also affected by the modalities that are primarily practiced in each month. For example, Shallow fishing has a high CPUE and August was the month with the highest E in Shallow fishing, so CPUE in August is very high. Likewise, for Razorfish fishing in September (Martorell-Barceló et al., 2021). Sociological phenomena could also be occurring. As an illustrative, speculative example, App users during summer months might be declaring longer E because they take a break from fishing (e.g. to eat or swim) and count this time as part of their fishing trip. In terms of the number of self-reports, July, August, September, October and December had the most, while March, April, May and June had the least. For E, significant differences were present in January, March and November (although the last two had only 13 Creel surveys each and might not be sufficient for proper statistical testing). For CPUE, no significant differences were associated to any month. Monthly, significant differences were few and are a good indication that App data is comparable to Creel survey data.

The most practiced modality in terms of E was Razorfish fishing, which has gained lots of popularity in the Balearic Islands (Alós et al., 2016). On the 1^st^ of September, hundreds of anglers go fishing for this species, but the fishery is short-lived as most of the E is concentrated during the first week (Signaroli et al., 2024). In this study, 82.03% of reported *Xyrichthys novacula* were captured in September. The species represented 25.56% of all captures in App data. Shallow fishing is the most popular modality in the Balearic Islands, practiced mainly during the morning by anglers of all levels of experience (March et al., 2014). This is reflected in both datasets in summatory of anglers per hour. In the Balearic Islands, its main captured species (i.e. *Serranus scriba, Diplodus annularis* and *Coris julis*) were reported to represent 61.2% of the recreational catch in terms of number of fish and 43.7% in terms of weight (Morales-Nin et al., 2005). In terms of number of fish in our study, limited to fishing inside MPAs, they represented 25.45%. Unsurprisingly, Razorfish fishing, Shallow and Deep fishing have the highest CPUE: Since in this study we calculated CPUE as the number of fish captured divided by E, rather than kg of fish divided by E, modalities that aim at smaller, easy to capture fish have the highest CPUE. Trolling fishing also accumulated a note-worthy summatory of E and was practiced mostly during 8:00 am to 14:00 pm. Self-reports show that this modality was practiced mainly in September and October, when the ban on *Seriola dumerili* juveniles and *Coryphaena hippurus* is lifted, and *Euthynnus alletteratus* juveniles and other little tunny species are abundant relatively near the coast (Navarro et al., 2017). For E, significant differences were present in Spearfishing, Shallow, Deep and Razorfish fishing comparisons. While Razorfish had few Creel surveys for a reliable comparison, Shallow and Deep fishing are the most sampled modalities in Creel survey and might present differences due to the misrepresentation of the fishing trip duration in both data collection methods, as explained above. Only Spearfishing presented significant differences among sources of data for CPUE, as well as E, showing that despite more hours of dedication, App users score more 0-catch or low-catch fishing trips. It would suggest that the level of experience of App users was lower or that these do not declare the totality of their harvest. Similar to a previous study (Riera-Batle & Grau, 2022), the most common captures of the modality were *Diplodus sargus, Labrus viridis, Mullus surmuletus, Octopus vulgaris, Sciaena umbra*a,nd *Scorpaena scrofa*.

In terms of the time of day when modalities are practiced, the datasets were highly similar with some displacement of peak activity time for Shallow, Razorfish fishing or Squid fishing. This displacement could be partly explained by mentioned Creel survey and App data limitations, like start-finish misdeclaration and unaccounted post-inspection E. It is important to note that not all Creel survey inspections collected time of start and finish, and this is especially true for Deep fishing inspections. Therefore, Deep fishing could not be properly compared, and neither Spinning, Razorfish and Bottom trolling fishing. With Razorfish fishing, since the species presents a wide range of awakening times and circadian rhythms (Akaarir et al., 2024; Martorell-Barceló et al., 2024), fish activity is spread over most daylight hours and so is E, unlike most of the other modalities which are mainly practiced in the morning. Squid fishing was mostly practiced at dusk and dawn, when the species is most active (Cabanellas-Reboredo et al., 2017).

When comparing MPAs it is important to consider the peculiarities of each one. Recent MPA like “Ses Bledes” and “Vedrà-Vedranell” still have few self-reports. Some are small and situated in less populated areas, like “Ses Bledes”, “Vedrà-Vedranell”, “Nord de Menorca” and “Illa de l’Aire”. On the other hand, “Migjorn de Mallorca”, “Badia de Palma” and “Illes del Toro i Malgrats” are bigger, close to yacht clubs and to very populated areas. The MPA “Migjorn de Mallorca” was of especial relevance as it accumulated the highest number of reported fishing trips (App - 36.47%, Creel survey – 40%) because it is the biggest MPAs and three major yacht clubs fall within its territory of 223.23 km^2^. E per trip was similar among a majority of MPAs and significant differences between App data and Creel survey data were limited to two MPAs: “Badia de Palma” and “Migjorn de Mallorca”. CPUE on the other hand was more variable among MPAs, and in this case less exploited areas like “Illa de l’Aire” or “Punta de sa Creu” yielded the highest CPUE. Significant differences among data sources for CPUE were present in “Badia de Palma” and “Freus d’Eivissa i Formentera” MPAs. The source of the mismatch between significantly different MPA remains unclear but the limited number of zones that coastguards can cover in vast MPAs may play a role. Regardless, the relative lack of significant differences among MPAs points to App data as a comparable source to Creel survey data for fisheries monitoring.

CPUE was calculated for some of the top captured species. Small, aggressive fish species captured by Shallow, Deep and Razorfish fishing modalities presented the highest CPUE. Trolling and Squid fishing species had lower values. The analysis did not explore in detail the exploitation of each species in terms of seasonality, fishing modality or location as it aimed to simply showcase the potential of the data, but these analyses could be performed for each of the 73 species for which there is data in App data, revealing relevant information for the monitoring and management of highly exploited and vulnerable species like, for example, *Epinephelus marginatus*or *Thunnus alalunga*. Significant differences between App data and Creel survey data in estimated species-specific CPUE were present in 4 out of 8 species. Creel survey was higher for *Coris julis*, *Diplodus vulgaris*and *Serranus scriba*, all of them the main captured species (notably missing *Diplodus annularis*). On the other hand, *Loligo vulgaris* was significantly higher in App data, although the average difference was very small (0.2 fish per fishing trip). Jiorle et al. (2016), comparing data from the app iAngler to the Marine Recreational Information Program survey of Florida, showed that median catch rate for particular species were roughly similar among data sources: *Centropomus undecimalis* (medians ranging from 0 to 2 in app and 1 to 2 in conventional survey), *Cynoscion nebulosus* (medians ranging from 1 to 3 in app and 1 to 4 in conventional survey), and *Sciaenops ocellatus* (medians ranging from 0 to 1 in app and 0 to 3 in conventional survey).As considered in Jiorle et al. (2016), median differences among data sources for species-specific CPUE are relatively small and App data may be considered useful for fisheries monitoring, even at the species level for particular species.

The MPAs in the Balearic Islands benefit from extensive control measures, including coastguard patrols, mandatory fishing permits, and the required declaration of fishing trip details. These measures enhance data collection reliability and reduce reporting biases, distinguishing the app from other similar data collection aproaches. When comparing data collected via the app to conventional creel surveys, some differences—particularly across fishing modalities—were observed, though the datasets were largely comparable. Despite its limitations, the app "Diari de Pesca Recreativa" offers several advantages: enforced compliance with declarations, a substantial volume of data (e.g., 953 self-reports in 2023), an absence of intrinsic modality bias, and a lack of temporal or seasonal biases commonly associated with creel surveys. Furthermore, app-based data collection is non-invasive, requires minimal time investment, and incurs low operational costs. Integrating data from both sources represents the most effective strategy, as they complement each other by compensating for respective limitations, thereby providing abundant, high-quality data for monitoring recreational fisheries in the MPAs of the Balearic Islands. Importantly, this approach opens opportunities for further innovation, such as using artificial intelligence (AI) to automate data acquisition from user-reported images. AI-driven image analysis could enhance the accuracy and efficiency of fish identification and automatization, and can provide size measurements, further maximizing the app’s potential for fisheries science and management. Developing tailored models to mitigate biases and fully harness this data source warrants further exploration.

## Acknowledgements

This work has been funded by the project (MEPRO 5220/2023. REMAR). Josep Alós received funding from the METARAOR Project (Grant num. PID2022-139349OB-I00) funded by MCIN/AEI/10.13039/501100011033/FEDER, UE. The authors would like to thank Eneko Aspillaga for his helpful input to this work, as well as all of the recreational fishers involved in the study.

